# Kinematics to Kinetics: Evaluating OpenGRF for Scalable Ground Reaction Force Estimation in Healthy and Pathological Gait

**DOI:** 10.64898/2026.06.03.729748

**Authors:** Muhammad Abdullah, Abdul Aziz Hulleck, Kinda Khalaf, Marwan El-Rich

## Abstract

Ground reaction forces (GRFs) are important markers of gait impairment and rehabilitation, but direct measurement with force plates is expensive and limited to laboratory settings. OpenGRF is an open-source framework that estimates GRFs from motion-capture data using musculoskeletal modeling, yet its validity in pathological gait remains uncertain. This study evaluated the accuracy of OpenGRF for estimating three-dimensional GRFs during gait in healthy adults and in people with Parkinson’s disease (PD), stroke, hip osteoarthritis (HOA), and total hip arthroplasty, and assessed its ability to reproduce clinically relevant GRF peak magnitudes and timings. OpenGRF estimates were compared with force-plate measurements in healthy participants and in cohorts with PD (OFF/ON medication), stroke, and HOA before surgery (M0) and 6 months after surgery (M6). Accuracy was quantified using root mean square error (RMSE), normalized RMSE (NRMSE), and Pearson correlation coefficients (PCC). Peak analyses examined biases in anterior-posterior (AP) braking and propulsive peaks, first and second vertical peaks (V1, V2), and the main mediolateral peak, as well as timing shifts across the gait cycle. One-dimensional statistical parametric mapping tested waveform differences (p = 0.01). OpenGRF reproduced overall GRF profiles across groups. Vertical GRF showed the best agreement (PCC 0.84-0.94; NRMSE 0.14-0.23), whereas mediolateral GRF showed low absolute error (RMSE 1.45-1.95 %BW) and moderate-to-good agreement (PCC 0.70-0.82). AP GRF was least accurate (PCC 0.61-0.73), especially during propulsion in PD (NRMSE up to 0.39). OpenGRF can reasonably estimate GRFs in healthy and pathological gait and may support kinetic gait assessment when force plates are unavailable.

## I. Introduction

Gait analysis is a fundamental quantitative tool for studying human locomotion, providing objective insights into neuromuscular control, biomechanical function, and compensatory strategies across a wide range of clinical populations [1]. Among biomechanical markers, ground reaction forces (GRFs) are key kinetic measures that characterize the interaction between the body and the ground during walking [2]. In particular, the vertical GRF (vGRF) is widely used to assess dynamic weight bearing, balance control, and overall gait performance [3]. In healthy gait, the vGRF typically exhibits an M-shaped pattern with two distinct peaks: during weight acceptance (V1) and during push-off (V2), separated by a mid-stance valley that reflects stabilization during single-limb support [4]. In the anterior-posterior (AP) direction, healthy GRF typically shows an initial negative braking phase followed by a positive propulsive phase, reflecting deceleration after heel contact and forward propulsion during late stance. In the mediolateral (ML) direction, GRF is smaller in magnitude and reflects side-to-side load transfer and dynamic balance control throughout stance [5]. Pathological gait refers to deviations from normal walking patterns caused by neurological, musculoskeletal (MSK), or systemic disorders, including stroke, Parkinson’s disease (PD), cerebral palsy, multiple sclerosis, peripheral neuropathy, and orthopedic conditions such as hip or knee osteoarthritis [5, 6].

GRF quantification is particularly valuable for characterizing disease-specific gait impairments. In PD, reduced peak propulsive forces and diminished push-off reflect bradykinesia and impaired motor control, while dopaminergic therapy such as Levodopa, the gold-standard pharmacological treatment, can partially restore peak propulsive forces and improve locomotor biomechanics [3, 6, 7]. After stroke, GRFs commonly reveal pronounced asymmetry between the paretic and non-paretic limbs, including decreased weight acceptance, reduced vertical loading on the affected side, and altered AP braking and propulsive forces [5, 8, 9]. In hip or knee osteoarthritis, GRF alterations reflect reduced loading on the painful limb and compensatory overloading of the contralateral limb, providing quantitative evidence of joint offloading strategies that may accelerate degeneration or contribute to secondary injury [10]. Similarly, after total hip arthroplasty (THA), although gait and pain often improve, GRF patterns frequently show persistent deficits, including reduced vertical loading on the operated limb, delayed peak loading, reduced propulsive AP forces, and interlimb asymmetry [11, 12]. Collectively, these findings highlight the importance of GRF magnitude and timing as objective biomarkers of gait impairment, functional capacity, and therapeutic response, supporting data-driven rehabilitation planning and outcome evaluation.

Force plates and instrumented walkways or treadmills remain the gold standard for GRF measurement, but their use has important technical and practical limitations. These systems require precise calibration, controlled laboratory conditions, and careful synchronization with motion capture (MoCap) systems, making data acquisition and analysis complex and time intensive. In addition, their high cost, large space requirements, and limited portability restrict their use in routine clinical practice and community settings [13]. Force-plate systems, in general, often capture only one valid step and require participants to target a specific foot placement area, which may disrupt natural gait and mask or exaggerate pathological abnormalities. Instrumented treadmills, meanwhile, typically assess only straight-line walking at constant speed, limiting their ecological validity. Together, these constraints underscore the need for continuous and more realistic gait monitoring approaches to support disease tracking and treatment evaluation [13].

Recent advances have aimed to overcome these limitations by enabling kinetic estimation outside controlled gait laboratories. One notable example is OpenGRF [14], an open-source framework that predicts GRFs directly from MoCap data using MSK modeling in OpenSim, thereby eliminating the need for force plates or instrumented treadmills during gait assessment. OpenGRF has shown strong agreement with experimentally measured GRFs, with normalized root mean square error (NRMSE) ranging from 1.07% to 7.84% across activities of daily living such as level walking and stair negotiation [14]. However, its evaluation has been largely restricted to healthy individuals, and its applicability in populations with pathological gait remains unclear. Given the well-documented GRF abnormalities in PD, hip osteoarthritis (HOA), THA, and post-stroke hemiplegia, it is important to determine whether OpenGRF can accurately reconstruct clinically meaningful GRF features, including weight acceptance, push-off, AP propulsive and braking forces, and their timing characteristics.

Therefore, this study addressed the following research questions: (1) How large is the overall error between OpenGRF-estimated and force plate-measured GRFs during gait? (2) What is the bias in clinically relevant GRF features, including the two vGRF peaks, AP propulsive and braking peaks, and maximum ML GRF, as well as their timing within the gait cycle? Answering these questions is essential to determine whether OpenGRF can support digital biomarker development, objective rehabilitation monitoring, and longitudinal assessment of disease progression outside traditional laboratory settings. Accordingly, this study aimed to assess the performance of OpenGRF in estimating GRFs during gait in healthy individuals, patients with PD, stroke survivors, and individuals evaluated before and after THA.

## II. Related work

In addition to MSK modeling-derived approaches, insole- and inertial measurement unit (IMU)-based systems have also been proposed in the literature to estimate GRFs in different directions. This section provides a brief overview of these methods.

### A. Musculoskeletal Modeling Approaches

To address the drawbacks of traditional force-plate systems, MSK modeling has emerged as alternative means for estimating GRFs by combining inverse dynamics and anatomical models. Recent studies report promising outcomes using such computational frameworks to estimate external kinetics without force-plate instrumentation using foot contact mechanism [14, 15]. Hulleck et al. [16] investigated the accuracy of GRF estimation using foot contact elements in the AnyBody Modeling System as compared to experimental force-plate measurements in stroke survivors. Their findings demonstrated good agreement between modeled and measured forces, highlighting MSK simulation as a cost-effective and clinically meaningful approach that enables gait analysis beyond restricted laboratory environments, particularly for neurological populations who struggle with constrained force-plate stepping.

### B. Insole Based Measurement Methods

Alternative approaches leverage wearable technologies, such as plantar pressure insoles and shoe-embedded load sensors, to estimate GRFs. Pressure-sensing insoles have been used to estimate vertical GRF and predict shear components using recurrent neural network models, notably long short-term memory (LSTM) sequence-to-sequence (Seq2Seq) architectures [17, 18]. Although moderately successful, these methods typically exhibit reduced performance for ML shear forces, are limited by small participant samples (n=16), and require specialized instrumentation worn inside footwear which also may alter natural gait and artificially mask or exaggerate pathological abnormalities [17]. Similarly, uniaxial load cells integrated in shoe soles have been used, in conjunction with LSTM-based prediction tools, to estimate three-dimensional GRFs from kinematic motion data in 81 subjects [18], with RMSE values of 65.12 N, 15.50 N, and 9.83 N for vertical, AP, and ML GRFs, respectively. However, restricting measurements to shoed conditions limits applicability to individuals with foot deformities or those unable to use instrumented footwear, including many clinical populations [18].

### C. IMU-Based Deep Learning Approaches

Mourot et al. [18] introduced a deep learning framework, leveraging IMUs to estimate vGRF and generate two-dimensional plantar pressure maps from joint kinematics based on a convolutional neural network. More recently, an optimized autoencoder-based Seq2Seq architecture demonstrated highly accurate prediction of GRF components from ankle position alone during level walking, reporting normalized RMSE (NRMSE) values below 0.15% for ML and AP GRF and up to 0.46% and 0.3% for left and vGRF, respectively [2]. Nevertheless, the model showed difficulty reconstructing peak vGRF and was validated only on healthy subjects, limiting its generalizability. Physics-informed neural networks have also been explored for vGRF estimation, demonstrating strong capability in replicating curve morphology. However, their robustness depends heavily on training diversity, requiring large heterogeneous datasets to avoid overfitting and enhance generalization across gait disorders [**Error! Bookmark not defined**.].

While a wide spectrum of methods has been proposed to overcome GRFs laboratory measurement restrictions, from MSK simulations to wearable sensors, artificial intelligence and deep learning, the literature reveals persistent challenges. These include limited generalizability to pathological gait, difficulty in reconstructing critical GRF features such as peak loading and propulsion, restricted instrumentation requirements, and dependence on controlled environments. Therefore, there remains a substantial need for accurate, unobtrusive, and scalable GRF estimation methods applicable to real-life gait and gait-impaired patient populations.

## III. Methods

The methodology comprises acquisition and pre-processing of experimental motion and force data, MSK modeling in OpenSim, estimation of GRFs from joint kinematics using OpenGRF, post-processing to segment, normalize, and resample gait cycles, and statistical comparison of measured and estimated waveforms. The workflow used to compare experimentally measured GRFs with OpenGRF estimations is summarized in Fig. 1.

**Fig. 1.**
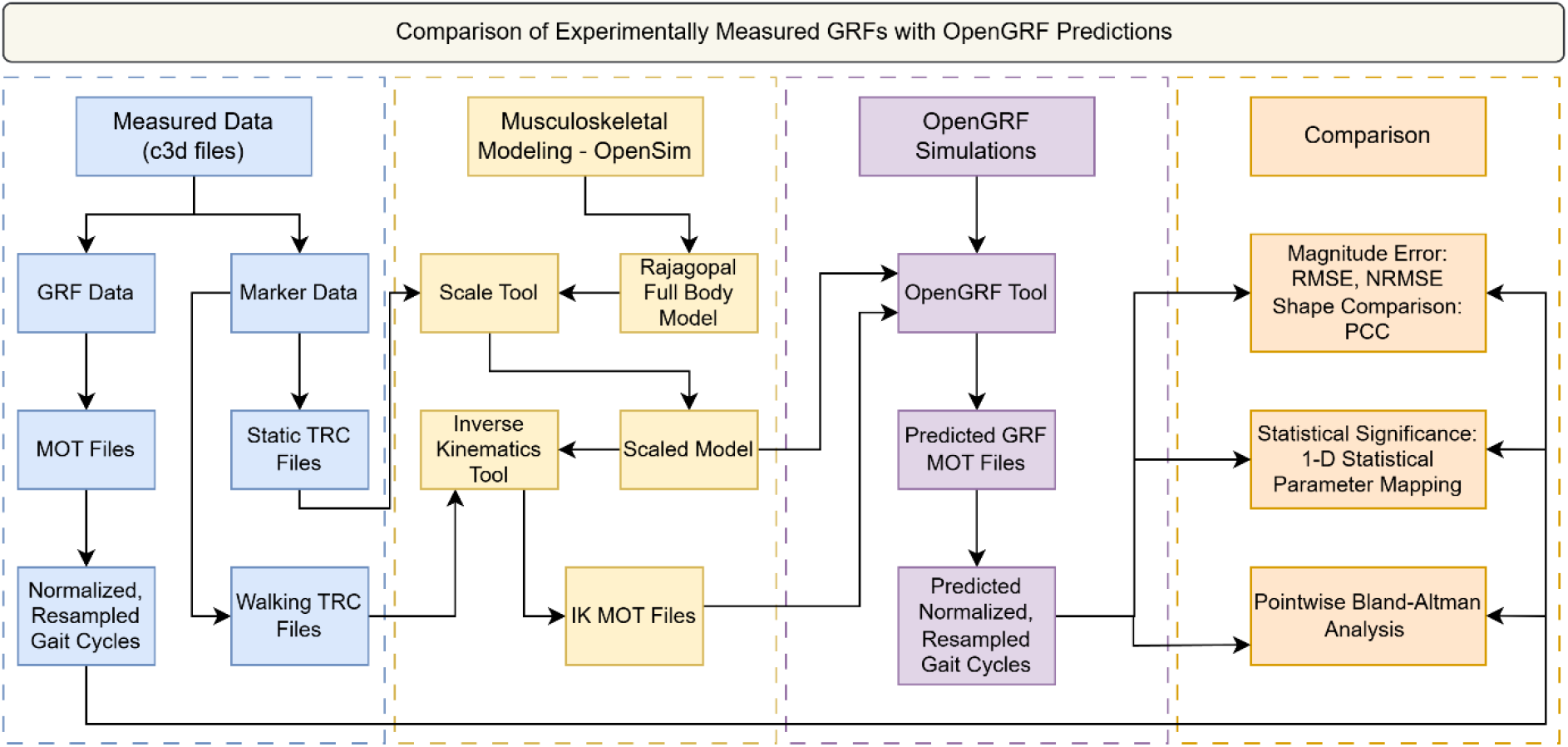
Schematic of the workflow used to compare experimentally measured ground reaction forces with OpenGRF estimations. Abbreviations: RMSE, root mean square error; NRMSE, normalized RMSE; PCC, Pearson correlation coefficient; c3d, coordinate three-dimensional file; MOT, motion data file; TRC, trace file.

**Fig. 1.**
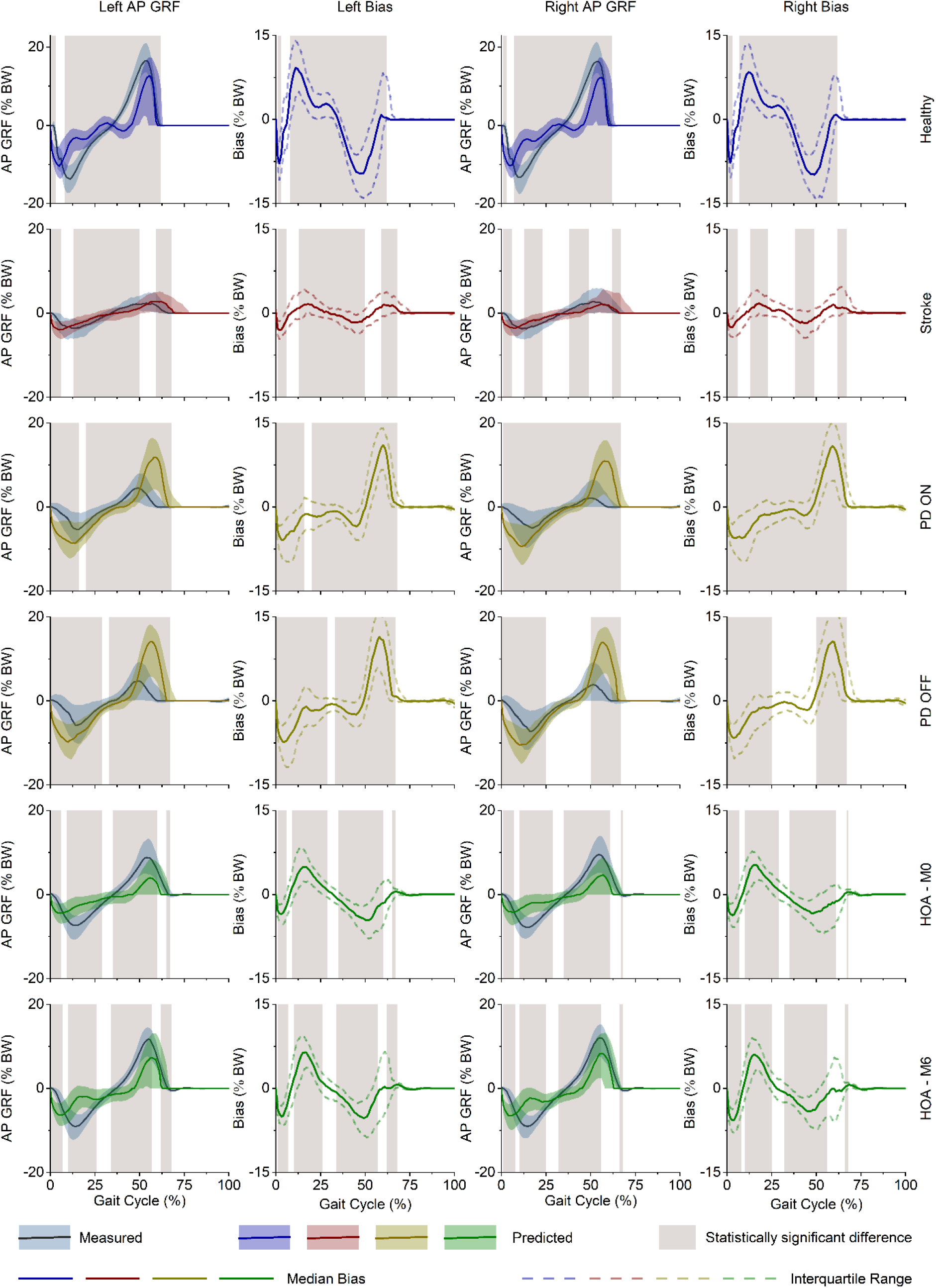
Comparison of measured and OpenGRF-estimated anterior-posterior ground reaction forces (AP GRF) across time-normalized gait cycles for the left and right limbs in healthy participants, stroke survivors, patients with Parkinson’s disease in the ON and OFF medication states, and individuals with hip osteoarthritis before surgery (M0) and 6 months after surgery (M6). Solid lines represent median waveforms, shaded bands indicate interquartile ranges (IQRs), and the bias plots show the median difference between estimated and measured signals with corresponding IQRs. Gray shaded regions denote gait-cycle intervals with statistically significant differences (p < 0.01) between measured and estimated AP GRFs.

### A. Measured Data and Subject Demographics

The motion data were obtained from three recent datasets comprising healthy controls, individuals with neurological disorders (stroke and PD), and individuals with MSK disorders (HOA). Participant demographic characteristics and details about MoCap equipment are summarized in Table I.

**TABLE I.** Demographics of participants and motion-capture system specifications.

| Parameter | Healthy | Stroke | PD | HOA-M0 | HOA-M6 |
| --- | --- | --- | --- | --- | --- |
| n | 138 | 50 | 26 | 106 | 92 |
| Age (years) | $51 \pm 20$ | $64 \pm 14$ | $64 \pm 10$ | $67 \pm 9$ | $68 \pm 9$ |
| Mass (kg) | $74 \pm 15$ | $72 \pm 14$ | $71 \pm 12$ | $78 \pm 17$ | $78 \pm 18$ |
| Height (mm) | $1684 \pm 103$ | $1705 \pm 80$ | $1660 \pm 70$ | $1642 \pm 88$ | $1645 \pm 87$ |
| Gender (m/f) | 65/73 | 34/16 | 20/6 | 51/55 | 44/48 |
| No of Cameras (Frequency) | 8 (100 Hz) | 8 (100 Hz) | 12 (150 Hz) | 8 (100 Hz) | 8 (100 Hz) |
| No of force plates (Frequency) | 3 (1000 Hz) | 3 (1000 Hz) | 5 (300 Hz) | 2 (1000 Hz) | 2 (1000 Hz) |
| No of Markers | 34 | 34 | 44 | 35 | 35 |
| Age (years) | $51 \pm 20$ | $64 \pm 14$ | $64 \pm 10$ | $67 \pm 9$ | $68 \pm 9$ |
| Mass (kg) | $74 \pm 15$ | $72 \pm 14$ | $71 \pm 12$ | $78 \pm 17$ | $78 \pm 18$ |
| Source | [19] | [19] | [20] | [21] | [21] |

#### 1) Healthy and Stroke Survivors

Motion data were obtained from Criekinge et al. [19] for healthy controls (n = 138) and stroke survivors (n = 50, 39 ischemic, 11 hemorrhagic). Stroke participants were 53 ± 19 days post-stroke, with lesions in the left hemisphere (n = 17) or right (n = 33). Their average functional ambulation category score was 3 ± 1. Data was captured using an eight-camera Vicon system (100 Hz) and four force plates (1000 Hz), tracking 14-mm markers based on the Plug-in Gait model. Each participant completed one static and three gait trials.

#### 2) Parkinson’s Disease

MoCap data for 26 individuals with idiopathic PD were obtained from Shida et al. [20]. Disease severity ranged from Hoehn & Yahr stage 1-4, and 13 had FoG. Each participant completed OFF- and ON-medicated sessions (≥12 h withdrawal vs. ~1 h post-dose). Data were collected using a 12-camera optical system synchronized with five force plates (GRF: 300 Hz; kinematics: 150 Hz) and 44 reflective markers. Each session included one static calibration and more than 10 overground gait trials.

#### 3) Hip Osteoarthritis and Total Hip Arthroplasty

Kinematic and kinetic data for unilateral HOA were obtained from Bertaux et al. [21] at two timepoints: preoperative M0 (30-1 days before surgery; n = 106: 43 left, 63 right) and postoperative M6 (~6-7 months; n = 92: 40 left, 52 right). Marker trajectories from 35 Plug-in Gait markers were recorded using eight Vicon MXT40 cameras (100 Hz) synchronized with two AMTI force plates (1000 Hz). Each session included one static calibration and more than 10 gait trials.

### B. Pre-Processing

MoCap data from all datasets were provided in the C3D file format. The preprocessing pipeline involved transformation of marker and force data to the OpenSim coordinate system, standardization and unit conversion of the signals, assignment and aggregation of force-plate data to the left and right limbs based on foot-marker proximity, and low-pass filtering of the force and moment signals using a 4th-order Butterworth filter with a 6 Hz cutoff frequency before exporting the processed trajectories and external loads into .trc and .mot files for OpenSim analysis. It should be noted that the static .trc marker file, generated from the static C3D trial, was used to scale the generic MSK model, whereas the gait .trc marker file, generated from the gait trial C3D, was used in inverse kinematics to estimate joint angles. The pre-processing steps were performed using custom-built MATLAB code.

### C. Musculoskeletal Modeling

MSK modeling was performed in OpenSim [22] using the full body Rajagopal model [23], with modifications reported by Lai et al. [24] and Uhlrich et al. [25]. The model included 22 body segments, 37 degrees-of-freedom, 17 joints, and 80 muscle-tendon units. Subject-specific personalization was performed using static .trc marker data and body mass with the OpenSim Scale Tool, producing a scaled model. Inverse kinematics was then applied to walking trials using the scaled model and .trc trajectories, minimizing marker error via weighted least squares, and joint angles were exported as .mot files for analysis.

### D. Estimation of GRFs - OpenGRF

OpenGRF [14] was used to predict GRFs for comparison with measured forces. It is an OpenSim based optimization workflow that estimates three-dimensional GRFs, ground reaction moments (GRMs), and center of pressure (CoP) directly from joint kinematics, given a subject specific scaled MSK model file (.osim) and a joint angle file (.mot). The tool returns time series of forces, moments, and CoP in OpenSim compatible .mot format. Further details on the OpenGRF tool can be found in [14]. It should be noted that, during simulation, the actuator optimal force magnitudes and weight factors were retained as originally defined in the tool, as these parameters could not be modified. During GRF estimation, kinematic tracking errors and pelvic residuals were monitored to ensure simulation accuracy. Summary results for the tracking errors and pelvic residuals are provided in supplementary information (SI)-A and B, respectively.

### E. Post-Processing

For each walking trial, two OpenSim .mot files were available, one containing measured GRFs and one containing OpenGRF estimations. Both forces were segmented into gait cycles for each limb, defined here as the stance phase from heel strike to toe off, using a vertical force threshold of 20 N. Heel strike was the first frame where the vertical component exceeded 20 N and toe off was the first frame where it fell below 20 N. Forces were then normalized to body weight and expressed as percent body weight (% BW). To enable point by point comparison, each cycle was time normalized to 0 to 100 percent and resampled to 101 samples.

In all cohorts, the estimated dataset contained more cycles than the measured dataset because force plates recorded only the steps that landed on the plates during a trial, whereas OpenGRF provided estimations for the entire gait trial. Because the number of cycles differed between measured and estimated forces, each trial and each limb were summarized by computing the median waveform across all available cycles. Then the analysis was restricted to trials for which both measured and estimated GRFs were available. Statistical comparisons on the resulting paired median waveforms were finally performed. The number of common gait cycles between the cohorts is reported in Table C.1 of SI-C.

Furthermore, for gait cycles matched between the measured and estimated datasets, peak magnitude and timing analyses were conducted using predefined gait-cycle windows to identify clinically relevant GRF features: AP-B (minimum AP force, 0-20% gait cycle), AP-P (maximum AP force, 40-65%), V1 (first vertical peak, maximum vertical force, 5-20%), V2 (second vertical peak, maximum vertical force, 35-60%), and ML-P (main ML extremum, defined as the maximum absolute ML force, 5-55%). Peak locations were determined independently for the measured and estimated waveforms.

### F. Statistical Analysis

Agreement between measured and estimated GRFs was quantified using magnitude and shape error metrics. For each GRF component, root mean square error (RMSE) and NRMSE were computed, with NRMSE defined as RMSE divided by the range of the measured waveform (maximum minus minimum). Waveform similarity was evaluated using the Pearson correlation coefficient (PCC) across the time-normalized gait cycle. The statistical significance of differences between the measured and estimated GRFs was assessed using one-dimensional Statistical Parameter Mapping (SPM) [26], with the significance level set at p = 0.01. Agreement was further characterized by pointwise Bland-Altman (BA) plots constructed across the gait cycle, in which the median bias (estimated minus measured) across trials was displayed together with non-parametric limits of agreement defined by the 25^th^ and 75^th^ percentiles. This combination of metrics captured absolute error, waveform concordance, and systematic bias.

Secondly, to quantify differences in GRF peak magnitude and timing, peak bias was defined as the difference between estimated and measured peak amplitudes, while timing error was defined as the difference in peak occurrence within the gait cycle. For each cohort, these outcomes were summarized across all matched gait cycles using the median and interquartile range (IQR). All analyses were performed using MATLAB R2025b.

## IV. Results

Subsequent subsections report the results for measured versus OpenGRF estimated GRFs across AP, ML, and vertical directions.

### A. Anteroposterior GRF

Figure 2 depicts the comparison between the measured and OpenGRF estimated AP GRF using time normalized gait cycles. Table II reports summary accuracy metrics, including median RMSE, NRMSE, and PCC with IQRs. For the AP GRF, OpenGRF generally reproduced the waveform shape in all groups. Absolute RMSE ranged from 2.28 to 5.75 % BW, with the lowest errors in stroke (2.28-2.38 % BW) and the highest in PD and healthy controls (up to 5.75 % BW in PD ON, right side). However, when normalized, the best AP accuracy was seen in healthy participants (NRMSE 0.17-0.18) and HOA (0.18-0.20), whereas PD showed the largest relative errors (NRMSE 0.32-0.39), especially on the right side in PD OFF. Correlations were moderate overall (PCC 0.61-0.71), with the highest PCC in HOA-M6 right (0.73) and the lowest in PD OFF right (0.61).

**Fig. 2.**
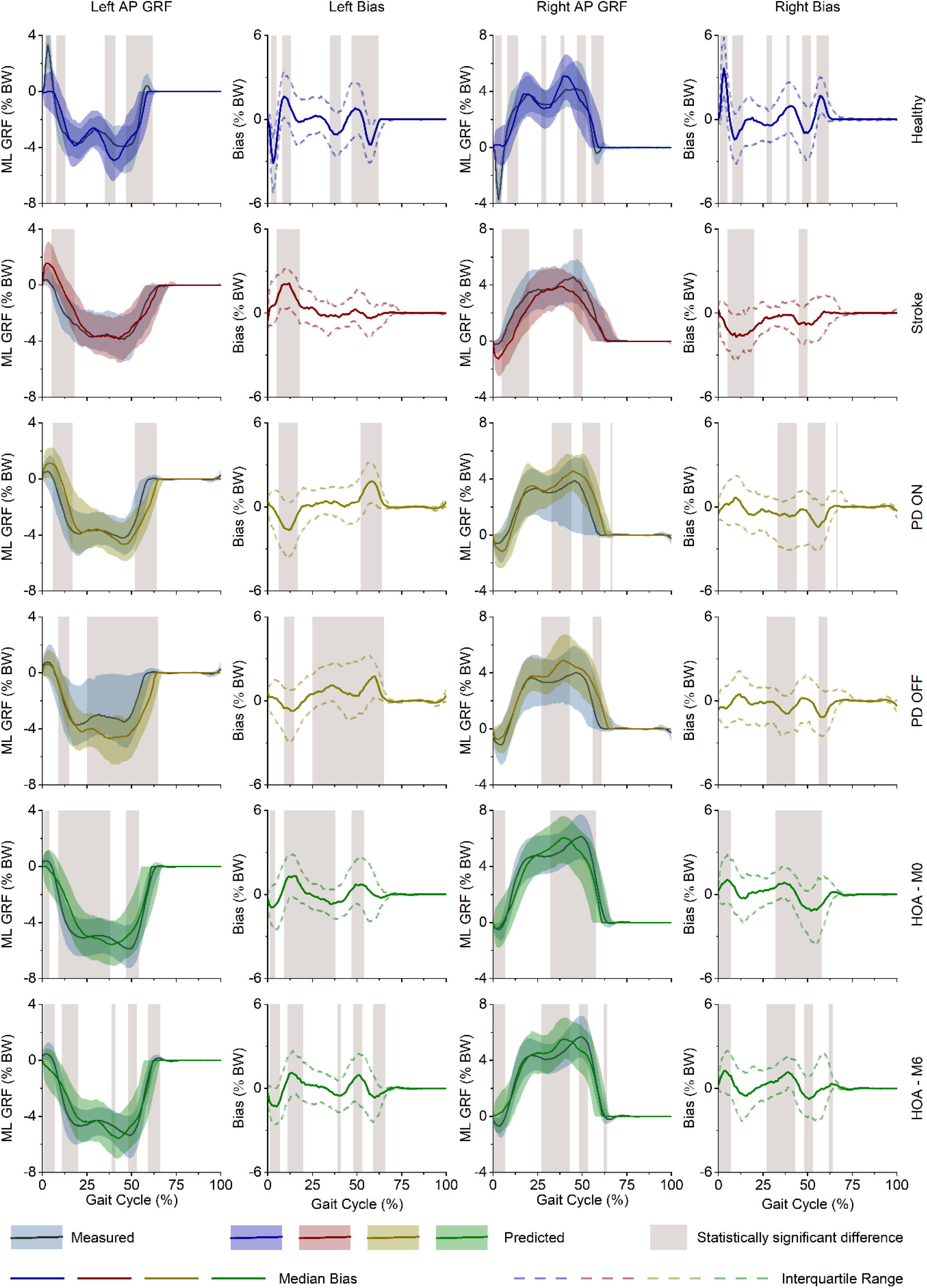
Comparison of measured and OpenGRF-estimated mediolateral (ML) ground reaction forces (GRFs) across time-normalized gait cycles for the left and right limbs in healthy participants, stroke survivors, patients with Parkinson’s disease in the ON and OFF medication states, and individuals with hip osteoarthritis before surgery (M0) and 6 months after surgery (M6). Solid lines represent median waveforms, shaded bands indicate interquartile ranges (IQRs), and the bias plots show the median difference between estimated and measured signals with corresponding IQRs. Gray shaded regions denote gait-cycle intervals with statistically significant differences (p < 0.01) between measured and estimated ML GRFs.

**TABLE II.**
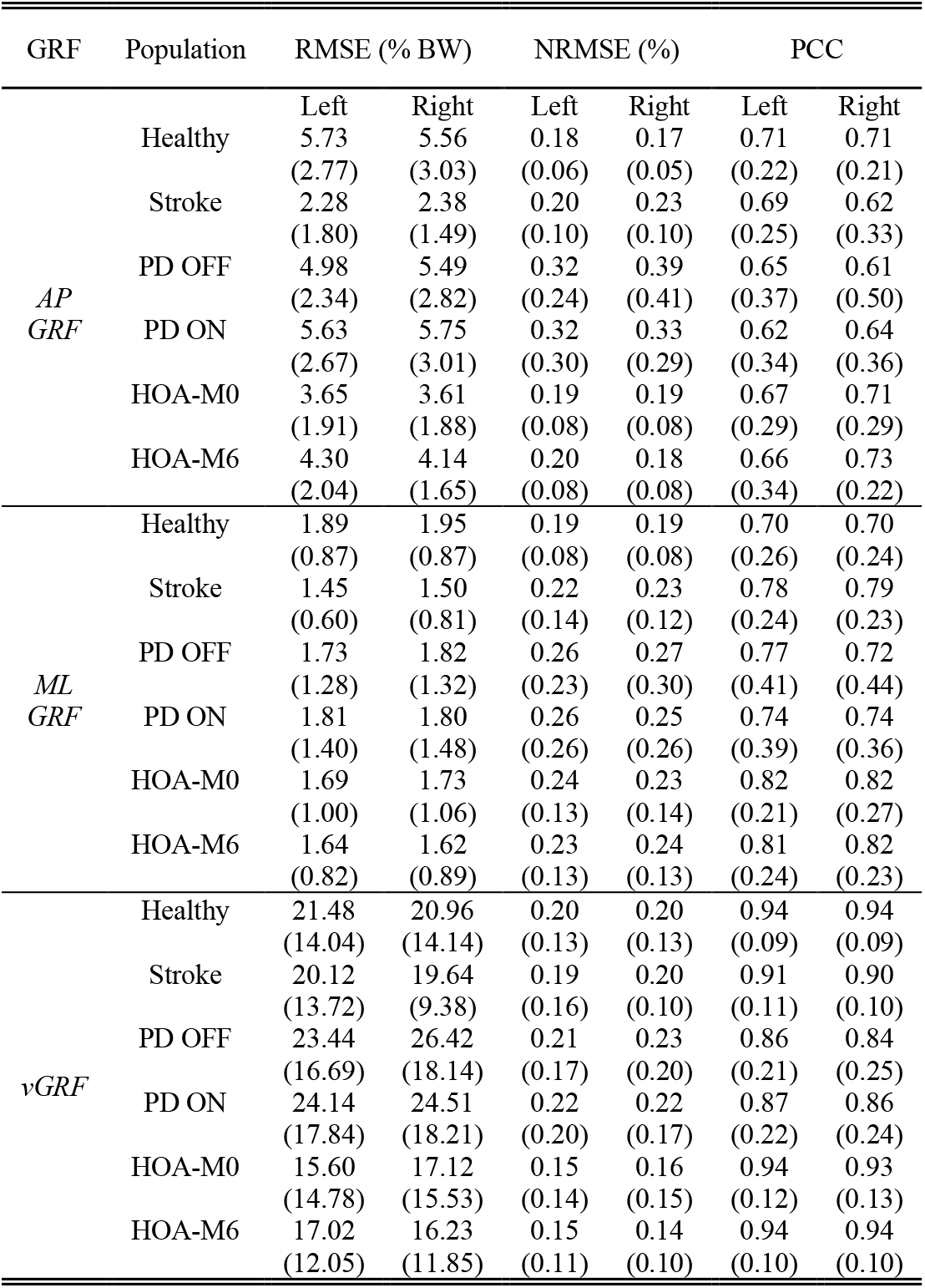
Median root mean square error (rmse), normalized rmse (nrmse), and pearson correlation coefficient (pcc) comparing measured and Opengrf predicted ground reaction force (grf) in three-dimensions. values in parentheses denote interquartile ranges.

From Figure 2, the main discrepancies were concentrated during the braking-to-propulsion transition and late stance push-off, where bias and statistically significant differences were most evident, particularly in healthy and PD groups. In contrast, stroke and HOA showed smaller AP amplitudes and comparatively lower absolute errors. Overall, AP GRF estimation was acceptable for capturing general trends, but propulsive-phase accuracy remains the main limitation, especially in PD.

### B. Mediolateral GRF

Figure 3 compares the measured and OpenGRF-estimated ML GRFs over time-normalized gait cycles, while Table II summarizes the median RMSE, NRMSE, and PCC, together with their IQRs. Overall, ML GRF estimation was more accurate than AP GRF estimation, with low absolute errors across all groups. Median RMSE ranged from 1.45 to 1.95 % BW, with the lowest values in stroke (1.45-1.50 % BW) and the highest in healthy participants (1.89-1.95 % BW). Relative errors were modest (NRMSE 0.19-0.27), being smallest in healthy controls (0.19 bilaterally) and largest in PD (0.25-0.27). Correlations were moderate-to-good overall, with the best agreement in HOA (PCC 0.81-0.82) and stroke (0.78-0.79), whereas healthy (0.70 bilaterally) and PD, particularly PD OFF right (0.72), showed comparatively lower agreement.

**Fig. 3.**
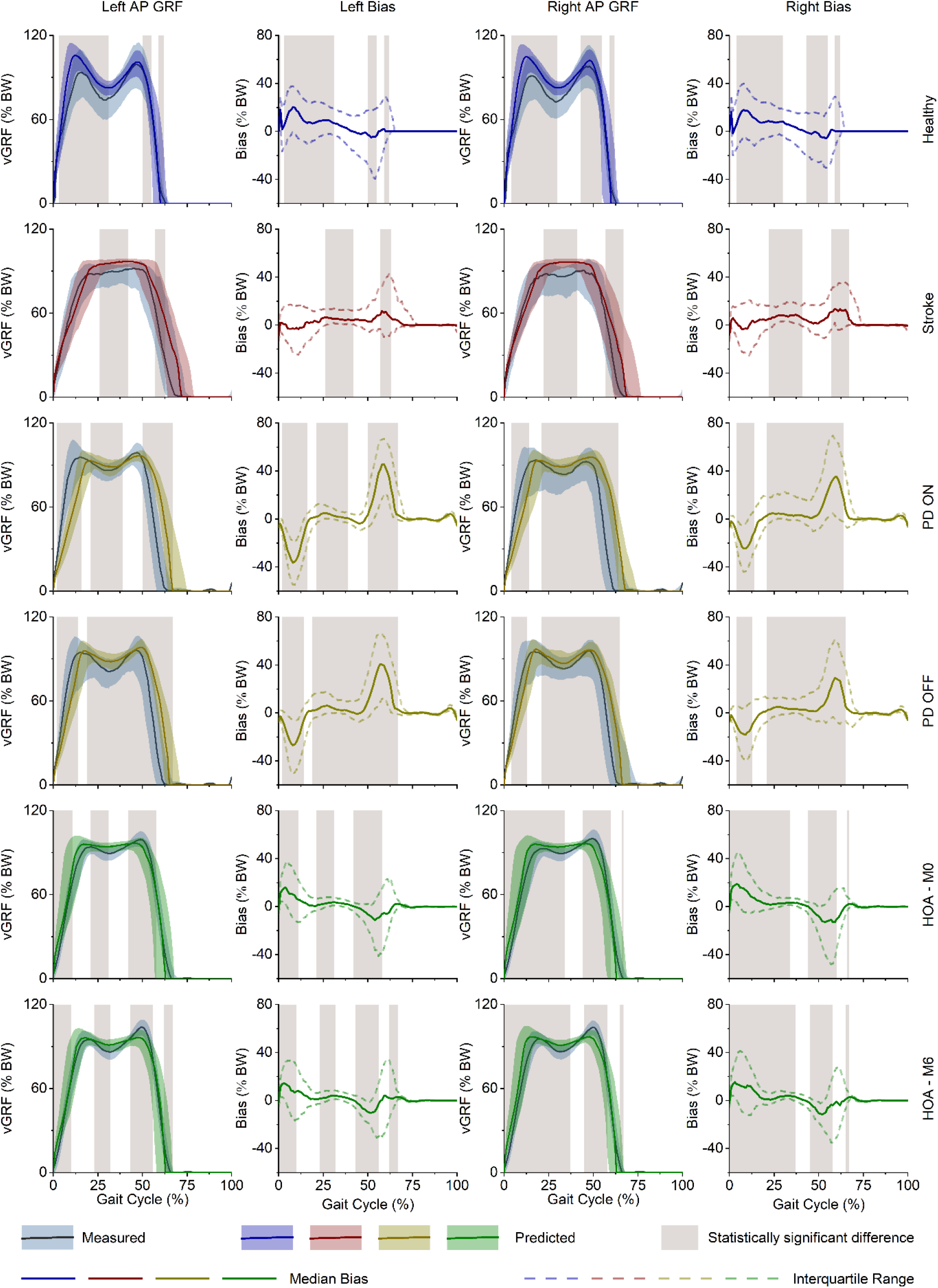
Comparison of measured and OpenGRF-estimated vertical ground reaction forces (vGRF) across time-normalized gait cycles for the left and right limbs in healthy participants, stroke survivors, patients with Parkinson’s disease in the ON and OFF medication states, and individuals with hip osteoarthritis before surgery (M0) and 6 months after surgery (M6). Solid lines represent median waveforms, shaded bands indicate interquartile ranges (IQRs), and the bias plots show the median difference between estimated and measured signals with corresponding IQRs. Gray shaded regions denote gait-cycle intervals with statistically significant differences (p < 0.01) between measured and estimated vGRFs.

From Figure 3, OpenGRF generally reproduced the overall ML waveform shape and side-to-side loading pattern, with small median bias around zero for most of stance. Statistically significant differences were intermittent rather than persistent, mainly during early stance and mid-to-late stance. Overall, ML forces were captured consistently across populations, with the strongest performance in stroke and HOA and slightly reduced accuracy in PD.

### C. Vertical GRF

Figure 4 presents the comparison between the measured and OpenGRF-estimated vGRF across time-normalized gait cycles, while Table II reports the median RMSE, NRMSE, and PCC, together with their IQRs. Overall, vGRF was the best-estimated GRF component across all populations, showing the strongest waveform agreement and highest correlations. Median RMSE ranged from 15.60 to 26.42 % BW, with the lowest errors in HOA (15.60-17.12 % BW) and the highest in PD, particularly PD OFF right (26.42 % BW). Relative errors remained low overall, with NRMSE between 0.14 and 0.23; the best performance was again observed in HOA-M6 right (0.14) and HOA-M0/M6 left (0.15), whereas the largest normalized errors occurred in PD OFF right (0.23) and PD ON/PD OFF left (0.22-0.21). Correlations were consistently high, ranging from 0.84 to 0.94, with the highest PCCs in healthy and HOA groups (0.93-0.94) and the lowest in PD OFF right (0.84).

From Figure 4, OpenGRF generally reproduced the characteristic double-peaked vGRF pattern well, with most discrepancies appearing around early weight acceptance, mid-stance, and late push-off, especially in PD. Overall, vGRF estimation was robust across groups, with best performance in healthy and HOA participants and slightly reduced accuracy in PD.

### D. Comparison of Peak GRFs Magnitude and Timing

Table III shows peak-specific magnitude bias and timing shift between measured and OpenGRF-estimated GRF features for the left and right limbs across all groups. Overall, vGRF peak timing was reproduced most consistently, with median time shifts generally close to 0 to ±4% of the gait cycle, whereas ML-P timing was more variable, as reflected by the wide IQRs. In terms of magnitude, ML-P showed the smallest bias across all populations (mostly within about ±1 % BW), indicating good agreement for ML-P amplitude.

**TABLE III.** Median (interquartile range) peak-specific bias and timing error between Opengrf-predicted and measured ground reaction force (grf) features for the left and right limbs in healthy participants and clinical cohorts with stroke, Parkinson’s disease (pd off and pd on), and hip osteoarthritis before surgery (hoa m0) and 6 months after surgery (hoa m6). Bias is expressed as the difference between predicted and measured peak magnitudes in % body weight (% bw), and time shift is expressed as the difference in peak occurrence in % gait cycle (% gc). Peak definitions are: ap-b, anterior-posterior braking peak; ap-p, anterior-posterior propulsive peak; v1, first vertical grf peak; v2, second vertical grf peak; ml-p, main mediolateral grf peak.

| Leg | Peak | Healthy |  | Stroke |  | PD OFF |  |
| --- | --- | --- | --- | --- | --- | --- | --- |
|  |  | Bias (% BW) | Time Shift (% GC) | Bias (% BW) | Time Shift (% GC) | Bias (% BW) | Time Shift (% GC) |
| *Left* | AP-B | 3.03 (9.02) | -5.00 (4.00) | -0.21 (3.75) | -4.00 (7.25) | -3.46 (4.21) | -1.00 (8.00) |
|  | AP-P | -3.72 (11.50) | 1.00 (5.00) | 0.83 (2.31) | 5.00 (7.25) | 6.57 (7.42) | 7.00 (8.00) |
|  | V1 | 14.99 (30.23) | -2.00 (5.00) | 5.89 (16.10) | 0.00 (8.00) | -5.33 (19.05) | 4.00 (7.00) |
|  | V2 | 2.86 (34.78) | 0.00 (5.00) | 4.56 (11.63) | 0.00 (8.25) | -2.30 (13.10) | 2.00 (7.00) |
|  | ML-P | -1.00 (3.31) | -1.00 (20.00) | -0.41 (1.93) | 0.00 (12.25) | 0.46 (2.63) | 1.00 (16.00) |
| *Right* | AP-B | 2.82 (9.19) | -5.00 (5.00) | 0.08 (3.47) | -4.00 (8.00) | -4.34 (5.99) | -2.00 (9.00) |
|  | AP-P | -3.61 (12.07) | 1.00 (5.00) | 0.98 (3.24) | 4.00 (8.00) | 8.71 (7.72) | 6.00 (10.00) |
|  | V1 | 16.51 (29.98) | -2.00 (6.00) | 4.95 (18.38) | 0.00 (5.00) | -1.03 (26.02) | 1.00 (7.00) |
|  | V2 | 2.40 (34.69) | 0.00 (6.00) | 5.06 (18.78) | -2.00 (7.00) | 0.72 (24.78) | 0.00 (9.00) |
|  | ML-P | 1.05 (3.20) | -1.00 (21.00) | 0.19 (1.90) | -2.00 (11.75) | -0.73 (3.20) | 1.00 (22.50) |
| Leg | Peak | PD ONN |  | HOA-M0 |  | HOA-M6 |  |
|  |  | Bias (% BW) | Time Shift (% GC) | Bias (% BW) | Time Shift (% GC) | Bias (% BW) | Time Shift (% GC) |
| *Left* | AP-B | -3.63 (5.89) | -2.00 (6.00) | 2.00 (4.21) | -6.00 (7.00) | 1.26 (4.64) | -6.50 (6.00) |
|  | AP-P | 7.83 (7.15) | 7.00 (9.00) | -1.45 (5.97) | 2.00 (8.00) | 0.20 (6.37) | 3.00 (6.50) |
|  | V1 | -1.94 (19.83) | 2.00 (7.00) | 3.75 (13.36) | -1.00 (5.00) | 3.92 (11.42) | -1.00 (5.00) |
|  | V2 | 0.29 (18.11) | 2.00 (6.00) | -1.70 (12.61) | -1.50 (8.00) | -5.10 (14.70) | -1.50 (8.00) |
|  | ML-P | 0.47 (3.13) | 5.00 (24.00) | -0.24 (1.98) | -1.00 (22.00) | -0.57 (2.60) | -1.50 (22.00) |
| *Right* | AP-B | -3.77 (5.66) | -2.00 (7.00) | 1.75 (4.27) | -7.00 (7.00) | 0.75 (4.37) | -8.00 (6.00) |
|  | AP-P | 8.93 (7.54) | 5.00 (9.25) | -1.14 (5.12) | 0.00 (8.00) | 0.08 (5.37) | 1.00 (7.00) |
|  | V1 | -0.17 (20.50) | 1.00 (7.00) | 5.07 (16.56) | -2.00 (7.00) | 4.55 (12.49) | -2.00 (6.00) |
|  | V2 | -0.66 (17.34) | 0.00 (9.75) | -1.65 (12.39) | -3.00 (8.00) | -6.46 (14.51) | -2.00 (7.00) |
|  | ML-P | -0.58 (2.81) | 2.00 (24.00) | 0.50 (1.84) | -7.00 (18.00) | 0.78 (2.22) | -4.00 (18.00) |

For the AP component, the largest bias was observed in PD, where AP-B was underestimated (about −3.5 to −4.3 % BW) and AP-P was overestimated (about 6.6 to 8.9 % BW) with a delayed occurrence of 5-7% GC, suggesting that push-off remains the most difficult AP feature to reconstruct. In contrast, stroke showed the smallest AP magnitude bias, with values close to zero for both braking and propulsion, although propulsion timing was still delayed by about 4-5% GC.

For vGRF, the largest bias was observed for V1 in healthy participants (about 15-17 % BW), while stroke, PD, and HOA generally showed smaller peak biases. In HOA, AP peak biases remained small at both M0 and M6, but braking peaks tended to occur earlier (about −6 to −8% GC) and V2 became slightly more underestimated at M6. Overall, the results suggest that OpenGRF captures peak magnitudes reasonably well for ML and most vGRF features, but AP propulsion is the least robust feature, particularly in PD.

## V. Discussion

This study evaluated the performance of OpenGRF for estimating three-dimensional GRFs from MoCap kinematics in healthy adults and clinical cohorts with PD, stroke, and HOA. Overall, the results showed that OpenGRF estimated the vGRF most accurately, with the highest waveform agreement across groups (PCC: 0.84-0.94) and low relative error (NRMSE: 0.14-0.23), despite larger absolute RMSE values due to the higher magnitude of the vertical force component. The ML GRF was also estimated with acceptable performance, showing low absolute errors (RMSE: 1.45-1.95 % BW), modest normalized error (NRMSE: 0.19-0.27), and moderate-to-good correlations (PCC: 0.70-0.82). In contrast, the AP GRF showed the lowest accuracy, with more variable agreement (PCC: 0.61-0.73) and larger relative errors in some groups, particularly in PD (NRMSE up to 0.39). Peak-based analysis further showed that AP propulsion was the least robust feature, with overestimation in PD of about 6.6-8.9 %BW and delays of 5-7% of the gait cycle. By comparison, vGRF peaks were reproduced more consistently, with timing shifts generally close to 0 to ±4% GC in most groups, although the first vertical peak (V1) showed the largest magnitude bias in healthy participants (about 15-17 %BW). The main ML peak showed the smallest magnitude bias overall, mostly within about ±1 %BW, but with wider timing variability across groups. OpenGRF performed best for vGRF because the vertical force is the main component of the gait loading pattern and is therefore easier to reconstruct. This agrees with the OpenGRF study [14], which also reported the highest accuracy for vertical GRFs and found the largest CoP errors during double support. In contrast, AP forces, especially propulsion, require more precise estimation of foot contact position and push-off timing, which likely made them harder to predict in this study. This effect was most evident in PD, where propulsion deficits were already more pronounced. The ML force was estimated with low bias overall, but its timing was less consistent, indicating that OpenGRF reproduced the size of the side-to-side force better than its exact timing. Taken together, these findings indicate that OpenGRF can generally recreate the overall three-dimensional GRF profiles across healthy and pathological gait and can additionally predict key clinically relevant peak features, especially the vertical GRF peaks and, with lower accuracy, the AP braking and propulsive peaks.

Hulleck et al. [16] used the AnyBody MSK model to predict GRFs for stroke survivors in the Van Criekinge et al. [19] dataset. Reported mean ± standard deviation RMSEs (% BW) were ML 5.08 ± 2.2 / 4.88 ± 2.2, AP 7.02 ± 4.4 / 5.28 ± 2.2, and vertical 44.0 ± 18.0 / 46.7 ± 23.0 for right/left legs. In this study, median ± IQR RMSEs (% BW), for stroke, were lower: ML 1.50 ± 0.81 / 1.45 ± 0.60, AP 2.38 ± 1.49 / 2.28 ± 1.80, and vertical 19.64 ± 9.38 / 20.12 ± 13.72 for right/left legs. OpenGRF outperformed the AnyBody implementation based on mathematical model by Fluit et al. [15]. These comparisons should be interpreted with caution, as the AnyBody models employed by Hulleck et al. [16] and the Rajagopal MSK model used in this study differ in their degrees of freedom, joint definitions, and muscle-tendon representations.

Eltoukhy et al. [27] used a Kinect-driven MSK model based on Fluit et al. [15] to predict GRFs in PD and reported mean errors of −1 % BW and −9 % BW for V1 and V2, and 3 % BW and −4 % BW for the AP shear maxima and minima respectively. In the present study, vGRF peak bias in PD was generally small, ranging from −5.33 to −0.17 % BW for V1 and −2.30 to 0.72 % BW for V2, whereas AP braking bias was comparable (−3.46 to −4.34 % BW) but AP propulsion was more strongly overestimated (6.57 to 8.93 % BW) and delayed by 5-7% GC. Thus, compared with Eltoukhy et al. [27], OpenGRF showed similarly good performance for vertical peaks and AP braking, but lower accuracy for AP propulsion. This comparison should be interpreted cautiously, as Eltoukhy et al. [27] studied a smaller PD cohort using Kinect-based input, whereas the present study used a larger sample and marker-based MoCap system.

This study has several limitations that warrant mention. First, the investigation was limited to pathological gait in stroke, PD, and HOA, and therefore did not consider other clinically relevant conditions such as cerebral palsy, knee osteoarthritis, or limb amputation. Second, the PD cohort was gender imbalanced, with only 6 of 26 participants being female, which constrains the generalizability of the findings. In addition, the Rajagopal MSK model used in this study is based on male anatomical geometry, and standard scaling procedures may not adequately capture gender-specific anatomical differences, potentially introducing bias in estimations for female participants. Furthermore, the analysis focused exclusively on GRFs, while CoP, free moment, and intersegmental joint forces and moments, derived from inverse dynamics, were not validated. Finally, MoCap data were acquired in a laboratory setting, which may not fully represent real-world walking or conditions encountered with wearable or marker-less sensing.

Future work should expand the range and diversity of clinical populations studied, improve gender balance, incorporate gender-specific MSK models, validate additional kinetic outputs (CoP, free moments, joint forces/moments), and evaluate performance under out-of-laboratory and wearable-sensor conditions.

Overall, while OpenGRF accurately reflects GRF waveform morphology across diverse populations, phase-specific errors indicate that further refinement, particularly of vertical force estimation and pathological gait dynamics, are warranted to improve clinical utility. Integration of improved foot-ground interaction models, subject-specific footwear and soft-tissue compliance parameters, and speed-dependent calibration may further support improved force estimation in future implementations.

## VI. Conclusion

This study evaluated the ability of OpenGRF to estimate three-dimensional GRFs from MoCap kinematics in healthy adults and in clinical populations with PD, stroke, and HOA before and after THA. Overall, OpenGRF was able to reconstruct the main GRF waveform profiles across all groups, supporting its potential as a practical alternative to direct force-plate measurements. Among the three components, vGRF showed the best agreement with measured data, with the highest waveform similarity and the most consistent reproduction of clinically relevant peak features. Similarly, ML GRF was also estimated with generally low bias, although peak timing was more variable. In contrast, AP GRF showed the lowest accuracy, particularly during late-stance propulsion, where larger magnitude and timing errors were observed, especially in PD. Peak-based analysis further confirmed that vGRF peaks and, to a lesser extent, AP braking and propulsive peaks can be reasonably estimated, although propulsion remains the least robust feature. Taken together, these findings indicate that OpenGRF can provide meaningful kinetic information in both healthy and pathological gait and may support digital biomarker development, rehabilitation monitoring, and longitudinal gait assessment beyond traditional laboratory-constrained settings.

## VIII. Author contributions

Conceptualization: MA and AAH

Data Curation: MA, and AAH

Investigation: MA and AAH

Methodology: MA, AAH, and ME

Supervision: ME and KK

Validation: MA and AAH

Visualization: MA

Writing – original draft: MA and AAH

Writing – review and editing: MA, AAH, KK, and ME

## IX. Ethical considerations

The motion data used in this study were obtained from publicly available datasets, for which participant consent had been obtained and reported by the original authors in the respective studies, as described in the Methodology section.

