## Supplementary Information for "Kinematics to Kinetics: Evaluating OpenGRF for Scalable Ground Reaction Force Estimation in Healthy and Pathological Gait"

### Predicting Ground Reaction Forces during gait in individuals with Parkinson's, Stroke, and OA diseases

#### SI-A: Tracking Errors

Table A.1 shows that inverse kinematics tracking errors were low across all cohorts and joints, indicating good agreement between the experimental marker data and the model-derived joint angles during GRF prediction. Overall, the largest median errors were observed at the ankle plantar/dorsiflexion and knee flexion/extension, particularly in the healthy and Hip OA groups, whereas the PD group showed the smallest errors across nearly all joints. Hip abduction/adduction had the lowest errors overall, while hip internal/external rotation and ankle plantar/dorsiflexion tended to show comparatively higher values. Errors were generally similar between the left and right limbs within each cohort. Taken together, these results suggest that kinematic tracking quality was consistently good and unlikely to be a major source of error in the GRF predictions.

**Table A.1.** Median (interquartile range) inverse kinematics (IK) tracking errors for lower-limb joint angles during GRF prediction in healthy participants and clinical cohorts with stroke, Parkinson's disease (PD), and hip osteoarthritis (Hip OA), reported separately for the left and right limbs. Abbreviations: Hip FE, hip flexion-extension; Hip AA, hip abduction-adduction; Hip IER, hip internal-external rotation; Knee FE, knee flexion-extension; Ankle PDF, ankle plantar-dorsiflexion.

| Joint Angle | Healthy | | Stroke | | PD | | Hip OA | |
|---|---|---|---|---|---|---|---|---|
| | Left | Right | Left | Right | Left | Right | Left | Right |
| Hip FE (°) | 0.56 (0.35) | 0.54 (0.36) | 0.37 (0.25) | 0.41 (0.29) | 0.30 (0.17) | 0.30 (0.20) | 0.46 (0.36) | 0.52 (0.36) |
| Hip AA (°) | 0.37 (0.26) | 0.36 (0.27) | 0.25 (0.18) | 0.28 (0.17) | 0.17 (0.12) | 0.16 (0.14) | 0.30 (0.21) | 0.29 (0.16) |
| Hip IER (°) | 0.99 (0.55) | 0.95 (0.60) | 0.59 (0.36) | 0.67 (0.47) | 0.28 (0.20) | 0.27 (0.19) | 0.72 (0.46) | 0.64 (0.53) |
| Knee FE (°) | 0.83 (0.62) | 0.82 (0.64) | 0.69 (0.53) | 0.70 (0.61) | 0.40 (0.46) | 0.44 (0.59) | 0.92 (0.75) | 0.97 (0.96) |
| Annkle PDF (°) | 1.24 (0.58) | 1.23 (0.48) | 0.73 (0.47) | 0.82 (0.64) | 0.48 (0.43) | 0.51 (0.40) | 1.07 (0.66) | 1.03 (0.66) |

#### SI-B: Pelvis Residuals

Table B.1 shows that the RMS pelvis residuals remained generally low across all cohorts, indicating good dynamic consistency of the simulations during GRF prediction. Residual forces in the AP, vertical, and ML directions were small overall, while residual moments for pelvic tilt, list, and rotation were also low in most groups. The largest median residual moment was observed for pelvic list in the Hip OA group (18.26 Nm), and the largest median residual force was observed for the vertical direction in Hip OA (4.76 N) and healthy participants (4.59 N). Nevertheless, these values remained well within the tolerance limits reported in the computed muscle control (CMC) online documentation (10 N for forces and 30 Nm for moments) [1]. Overall, these findings suggest that the residual loads required to drive the simulations were small and that the predicted motions were dynamically acceptable across all cohorts.

**Table B.1.** Median (interquartile range) root mean square (RMS) pelvis residual forces and moments during GRF prediction in healthy participants and clinical cohorts with stroke, Parkinson's disease (PD), and hip osteoarthritis (Hip OA). Residual moments are reported for pelvic tilt, list, and rotation (Nm), and residual forces are reported in the anterior-posterior (AP), vertical, and mediolateral (ML) directions (N). All values remained well within the tolerance limits reported in the computed muscle control (CMC) online documentation (10 N for forces and 30 Nm for moments).

| RMS Pelvis Residuals | Healthy | Stroke | PD | Hip OA |
|---|---|---|---|---|
| Tilt (Nm) | 14.77 (10.41) | 0.63 (2.22) | 0.22 (5.27) | 1.63 (4.00) |
| List (Nm) | 0.54 (4.32) | 0.00 (0.01) | 0.00 (0.35) | 18.26 (8.70) |
| Rotation (Nm) | 0.37 (2.09) | 0.00 (0.00) | 0.00 (0.11) | 3.77 (7.22) |
| AP (N) | 1.49 (6.19) | 0.75 (2.32) | 0.19 (5.02) | 1.23 (5.93) |
| Vertical (N) | 4.59 (1.82) | 0.08 (0.66) | 0.03 (30.31) | 4.76 (6.12) |
| ML (N) | 1.50 (7.35) | 0.00 (0.01) | 0.00 (1.14) | 1.07 (7.57) |

#### SI-C: Summary of Gait Cycles and Total Common Gait Trials

In all cohorts, the estimated dataset contained more cycles than the measured dataset because force plates recorded only the steps that landed on the plates during a trial, whereas OpenGRF provided estimations for the entire gait trial. Because the number of cycles differed between measured and estimated forces, each trial and each limb were summarized by computing the median waveform across all available cycles. Then the analysis was restricted to trials for which both measured and estimated GRFs were available. Statistical comparisons on the resulting paired median waveforms were finally performed. The number of common gait cycles between the cohorts is reported in Table C.1.

**Table C.1:** Summary of the total number of gait cycles common between the measured and OpenGRF predicted for three-dimensional ground reaction forces. Abbreviations: PD, Parkinson's disease (PD); HOA, hip osteoarthritis; M0, preoperative (before total hip arthroplasty); M6, 6 months post-total hip arthroplasty

| Population | Total Number of Gait Cycles | |
|---|---|---|
| | Left | Right |
| Healthy | 366 | 374 |
| Stroke | 61 | 58 |
| PD OFF | 345 | 313 |
| PD ON | 343 | 357 |
| HOA - M0 | 284 | 286 |
| HOA - M6 | 292 | 278 |
